# Assessing Immune Factors in Maternal Milk and Paired Infant Plasma Antibody Binding to Human Rhinoviruses

**DOI:** 10.1101/2023.12.17.565204

**Authors:** Jessica M. Vera, Sean J. McIlwain, Samantha Fye, Ann Palmenberg, Yury Bochkov, Hanying Li, Richard Pinapati, John Tan, James E. Gern, Christine Seroogy, Irene M. Ong

## Abstract

Before they can produce their own antibodies, newborns are protected from infections by transplacental transfer of maternal IgG antibodies and after birth through breast milk IgA antibodies. Rhinovirus (RV) infections are extremely common in early childhood, and while RV infections often result in only mild upper respiratory illnesses, they can also cause severe lower respiratory illnesses such as bronchiolitis and pneumonia. We used high-density peptide arrays to profile infant and maternal antibody reactivity to capsid and full proteome sequences of three human RVs - A16, B52, and C11. Numerous plasma IgG and breast milk IgA RV epitopes were identified that localized to regions of the RV capsid surface and interior, and also to several non-structural proteins. While most epitopes were bound by both IgG and IgA, there were several instances where isotype-specific and RV-specific binding were observed. We also profiled 62 unique RV-C dominant protein loop sequences characteristic of this species’ capsid VP1 protein. Many of these RV-C sites were highly bound by IgG from one-year-old infants, indicating recent or ongoing active infections, or alternatively, a level of cross-reactivity among homologous RV-C sites.

## Introduction

Maternally derived antibodies are an important form of humoral immunity for both the developing fetus and postnatal infant. Breast milk (BM) is replete with immunological and bioactive molecules that can supplement and stimulate the developing infant immune system (Lawrence and Pane 2007; Chirico et al. 2008; Kim 2021). Viral respiratory infection is the leading cause of hospitalization and acute health care costs for children under the age of five years (Witt, Weiss, and Elixhauser 2006; Duan et al. 2023). However, breastfeeding reduces the risk of upper respiratory infections during infancy (López-Alarcón, Villalpando, and Fajardo 1997; Bachrach, Schwarz, and Bachrach 2003; Jenkins et al. 2004; Sadeharju et al. 2007). While the mechanisms of protection conferred by breastmilk (BM) remain largely uncharacterized, IgG and IgA antibodies have been directly implicated in protection against two common respiratory viruses, respiratory syncytial virus (RSV) and human rhinovirus (RV), respectively (May and Clarke 2000; Mazur et al. 2018).

RSV and RV infections in children commonly cause mild upper respiratory illnesses but can also cause severe lower respiratory illnesses such as bronchiolitis and pneumonia. Historically, RSV has received more attention for its predilection for causing bronchiolitis in infants. However, RV-A and RV-C species are the most frequent viruses detected in children over one year of age who have wheezing illnesses or exacerbations of asthma (Miller et al. 2007; Lau et al. 2009; Lee et al. 2012; Han, Rajput, and Hershenson 2019; Biagi et al. 2020). These findings have spurred interest in characterizing infant and maternal humoral immunity to RVs. While the neutralizing immunogenicity of localized RV-A and RV-B capsid surface sites has been well demonstrated, a complete proteome-wide characterization of regions of RV bound by antibodies, including non-structural proteins, is lacking. Additionally, the overlapping contributions of BM relative to induced infant anti-RV antibodies to overall immunity are unknown.

We and others have developed methods for viral humoral profiling using peptide arrays that allow for expansive examination of antibody reactivity to short, linear peptide fragments (Asano et al. 2011; Kuivanen et al. 2014; Stephenson et al. 2015; Sabalza et al. 2017; Zhang et al. 2018; Heffron et al. 2018; Farrera-Soler et al. 2020; Heffron et al. 2021). To better understand how placental transfer and human BM confers protection against viral infections and affects infant immune development, we profiled antibody reactivity from BM, cord blood, and infant plasma samples collected from our ongoing observational birth cohort studies. Both IgA and IgG reactivity was profiled against overlapping linear peptide probes covering the capsid or proteomes of three commonly circulating RV genotypes (A16, B52, and C11) and for 62 putative RV-C-specific VP1 capsid antigens using high-density peptide arrays.

## Material and Methods

### Human subjects

The participants are from an ongoing prospective observational birth cohort study in rural Wisconsin designed to investigate early life farm exposure influence on immune development and respiratory allergies and illnesses in children (Seroogy et al. 2019). The study was approved by the institutional review boards at the Marshfield Clinic and the University of Wisconsin, and all participants provided informed consent before participating in the study. Infant participants in the ongoing study were born between 2013-2020. Infant participants born between 2013-Jan 2018 were enrolled from rural communities and farms in the vicinity of Marshfield, WI. Starting in February 2018, farm, non-farm, and traditional agrarian families were recruited concurrently at two study sites, the Marshfield Clinic and the LaFarge Medical Clinic (part of Vernon Memorial Healthcare). Written informed consent was obtained by the mother. Study procedures for enrolled participants include questionnaires, environmental assessments, and samples of blood and breast milk using standardized collection kits and procedures (Seroogy et al. 2019).

### Peptide microarray design and synthesis

The synthesized arrays included sequences covering the complete proteomes of human rhinoviruses B52, and C11, the capsid polyprotein sequence of RV A16, and 62 RV-C “C-loop” sequences (Tapparel et al. 2009; Nakagome et al. 2014; Gern et al. 2019). Peptides covering two exogenous viral protein sequences (Epstein-Barr virus (EBV) nuclear antigen 1 and cytomegalovirus (CMV) glycoprotein B) were included as controls. All proteins were tiled as 16 amino acid peptides, in register, overlapping by 15 amino acids. Additional peptides, 15-12 amino acids in length, were tiled at the protein C-terminus to increase coverage of these residues.

The chosen viral peptide sequences were printed onto peptide microarrays (Nimble Therapeutics Madison, Wisconsin, USA) (Heffron et al. 2018) using a Nimble Therapeutics Maskless Array Synthesizer (MAS) and light-directed solid-phase peptide synthesis using an amino-functionalized support (Geiner Bio-One) coupled with a 6-aminohexanoic acid linker and amino acid derivatives carrying a photosensitive 2-(2-nitrophenyl) propyloxycarbonyl (NPPOC) protection group (Orgentis Chemicals). Peptides with potential sequence overlap were placed in non-adjacent positions on the array to minimize impact of positional bias. Each array consisted of 12 subarrays containing up to 391,551 unique peptide sequences to probe each sample.

### Peptide microarray sample assay

Biospecimens were uniformly collected using centralized collection kits and standardized procedures. Plasma samples were obtained from blood collected in sodium heparin tubes, centrifuged for separation of cells and plasma, and plasma was removed and stored at -80°C until use.

Breast milk (BM) was collected at infant’s 2-month study visit. BM samples were thawed on ice and centrifuged (10,000 x g for 10 minutes at 4 °C), to separate fat and cells from whey. The fat layer was carefully removed with a sterile spatula and any residual lipids were aspirated using a wide-bore pipette tip. The clear supernatant containing whey and plasma samples were stored at -80°C until use.

IgG binding was assayed as previously described (Heffron et al. 2021). IgA primary sample binding was detected via FITC-conjugated goat anti-human IgA (Jackson ImmunoResearch; RRID: AB_2337652), which was scanned at 488 nm.

### Peptide microarray data analysis and visualization

pepMeld (https://github.com/DABAKER165/pepMeld) was used to process the raw array fluorescence intensity data as previously described (Heffron et al. 2021). Raw fluorescence signal intensity values were log_2_ transformed, and clusters of fluorescence intensity of statistically unlikely magnitude, indicating array defects, were identified, and removed. Local and large area spatial corrections were then applied, and the median transformed intensity of peptide replicates was determined. The resulting data was cross normalized using quantile normalization. Lastly, probe fluorescence intensity values were smoothed by averaging the signal intensities across a three-probe window centered on each probe. For each sample type, two technical replicates for at least two separate samples were profiled. A Pearson correlation threshold of ≥ 0.73 was used to filter pairs of technical replicates to allow most technical replicates to be retained. The smoothed signal for remaining technical replicates were averaged for each sample before any calls were made concerning epitope probabilities.

Heatmaps representing the array data were created using the ComplexHeatmap version 2.6.2 (Gu, Eils, and Schlesner 2016) package in R.

### Statistical analysis

Each test sample was profiled for both IgG and IgA reactivity. However, human BM contains predominantly secretory IgA while plasma has much more IgG (Hurley and Theil 2011). The observed differences in the antibody reactivity profiles between BM IgA and plasma IgG were consistent with reported differences in the abundances of these two immunoglobulins in these fluids. In addition, BM IgA displayed higher overall signal intensity and lower signal-to-noise ratio compared to plasma IgG. These differences could be due to the compositional differences between plasma and BM, specificity in anti-IgG and anti-IgA secondary antibodies used in our assay, or the overall specific activity of IgA and IgG antibodies. As a result, we processed plasma IgG data and BM IgA data as separate sample populations and defined sample-type specific epitope calling criteria for each.

Sample population mean and standard deviation were used to calculate a z-score for each probe. A z-score threshold of 3 for plasma IgG and 2.5 for BM IgA samples were set to define those probes with statistically significant antibody reactivity. “Epitopes” were defined as consecutive runs sequence-adjacent probes scoring above the z-score threshold in each test sample. Non-consecutive, singleton probes with significant reactivity in only a single test sample were discarded. Statistical analyses were performed in R version 4 using in-house scripts (see Data and code availability).

### Protein structures

The virion capsid structures of A16 (1aym), B14 (4rhv), and C15 (5k0u) have been determined to atomic resolution. Residue coordinates were obtained from https://rcsb.org. The array-profiled sequences of B52 and C11 are direct homologs to the capsid proteins of B14 and C15 respectively, so the analogous peptides (B52, C11) could be readily identified by sequence alignment and annotated on the known structures (B14, C15). Within the PyMol coordinate display program, the data2bfactor Python script (The PyMOL Molecular Graphics System, Version 2.0 Schrödinger, LLC) was used to substitute numerical parameters representing the peptide array data, replacing each coordinate file’s B-factor column.

### RV molecular detection and typing

Nasal swabs collected from participating infants at the indicated surveillance timepoints and during illness episodes were assayed for all common respiratory viruses by multiplex reverse transcription polymerase chain reaction (NxTAG Respiratory Pathogen Panel, Luminex, Austin TX). RV isolates were partially sequenced to identify the species and type and to differentiate lengthy single infections from serial infections with multiple RV strains (Bochkov et al. 2014).

### RV Capsid Protein Alignments

All RV-A and RV-B share strong capsid structure conservation and readily identifiable sequence homology in key internal protein sub features, making it possible to accurately predict and identify analogous NIm surface locations for equivalent viruses. Analogy-based RV capsid protein sequence alignments were generated as previously described (Palmenberg et al. 2009).

### RV-C VP1 C-loops homologous sequence clusters

Refined C loop multiple sequence alignment and Neighbor-Joining (NJ) tree were created using MUSCLE version 3.8.31 (Edgar 2004). Using the R phylogram package (v 2.1.0) (P. Wilkinson and K. Davy 2018) the NJ tree was processed to allow conversion to a hierarchical cluster object and cut into 12 clusters.

### Data and Code Availability

All peptide array datasets and code used will be made available after publication.

## Results

### Controls for antibody profiling

A newborn’s capacity to mount adaptive immune responses is limited and largely depends on innate and passive immunity for protection against infectious disease. Infant passive immunity is conferred *in utero* via placental transfer of maternal peripheral blood IgG antibodies, and by IgA and IgG antibodies acquired postnatally via breast milk (BM). By one year of age, transplacental antibodies have typically waned and the maturing infant immune system is the sole source of antibodies, induced thereafter by acquired infections and vaccinations. The landscape of maternally contributed and infant-derived antibodies to three species of RV were profiled for both IgG and IgA reactivity from 275 participant samples, including BM, cord blood, and infant plasma at 1 year of life (**Table 1**). The cohort was partially concordant, with a total of 54 sets of BM, cord blood, and infant plasma samples from the same mother-infant dyads.

**Table 1.**

| Sample | Count | Collection Time Point | Maternal Age (years)<sup>a</sup> | Humoral Profile | Antibody |
| --- | --- | --- | --- | --- | --- |
| breast milk | 108 | 2 months | 30.4 ± 4.2 | maternal | IgA |
| cord blood plasma | 93 | birth | 30.7 ± 4.3 | maternal | IgG |
| infant plasma | 74 | 1 year | 31.4 ± 3.9 | infant | IgG |

<sup>a</sup>mean age of mother at child's birth ± one standard deviation

In addition to testing for IgG and IgA reactivity to RV, we also profiled for reactivity to viral antigens from Epstein-Barr virus (EBV) nuclear antigen 1 (P03211) and cytomegalovirus (CMV) glycoprotein B (P06473) (**Fig. 1**). Nearly all adults have antibody reactivity to at least one of these viruses so their antigens provided powerful positive controls, especially for BM (Rumpold et al. 1987; Dillner et al. 1984; Meyer, Masuho, and Mach 1990; Cheng et al. 1991; Kniess et al. 1991; Ruprecht et al. 2014). Indeed, we found all tested BM IgA and cord blood IgG samples showed significant reactivity to at least one of these probes. However, a smaller proportion of infant samples showed similar strong reactivity to these controls when compared to maternal BM and cord blood. Individual infant samples also showed notable variability in specific epitope binding and the signal intensities. For example, infant EBV IgG reactivities were largely confined to the alanine-glycine rich repeats of nuclear antigen 1 (probes ∼87-325) and moreover had lower signal intensity compared to maternal (mother-child dyad) cord blood IgG (**Fig. S1A**). The IgG reactivity to EBV nuclear antigen 1 peptides among infant samples moderately correlated with cord blood IgG reactivity (median Pearson’s *r* = 0.66, N = 61).

**Figure 1.**
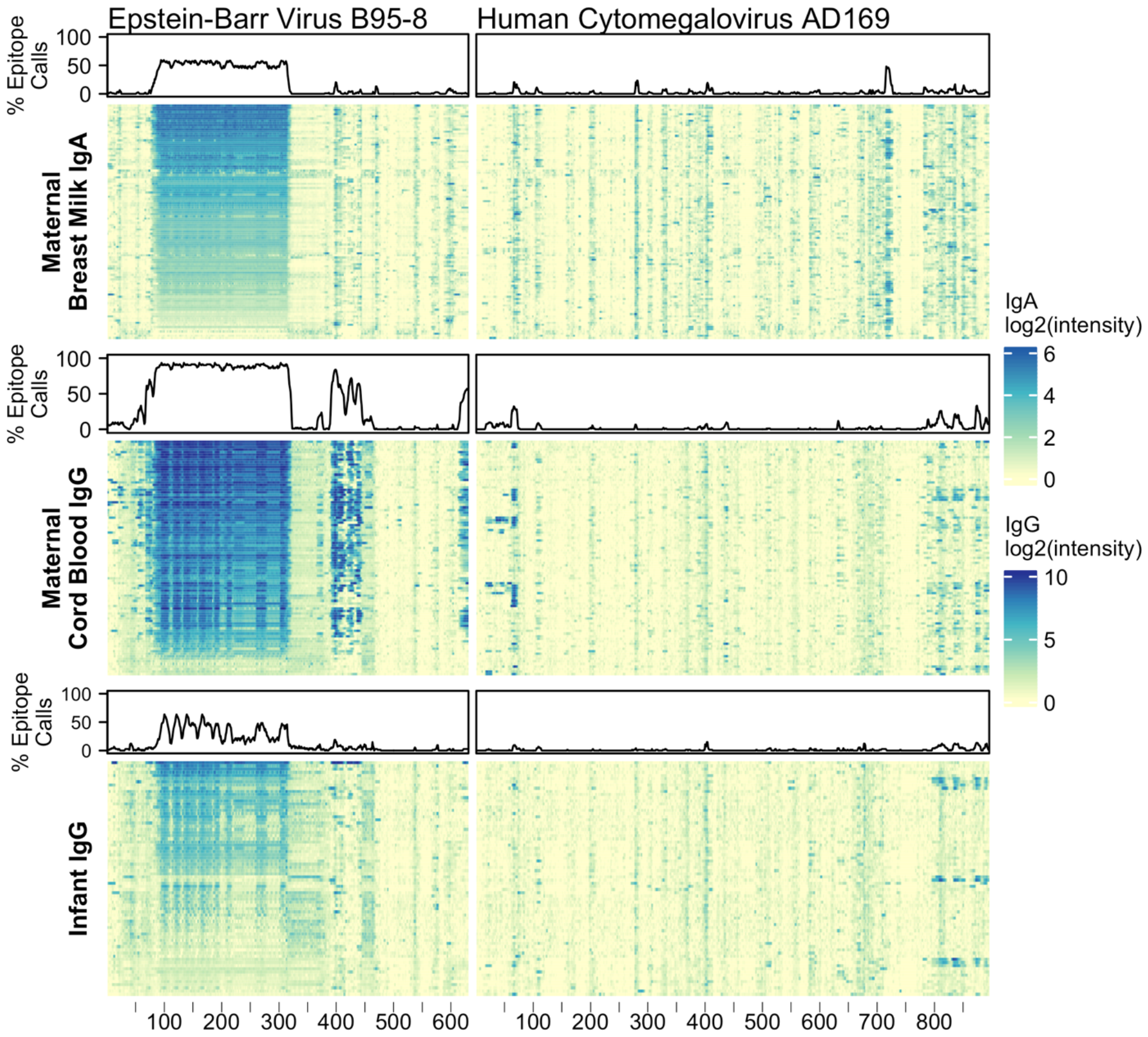
Maternal IgA and IgG and infant IgG reactivity to common viral antigen control proteins. Heatmap of antibody reactivity to tiled Epstein-Barr Virus nuclear antigen 1 (P03211) and Cytomegalovirus glycoprotein B (P06473) peptide sequences on a peptide microarray. IgG and IgA sample data were processed as distinct populations. Above each sample population is a line plot indicating the percent of samples with statistically significant reactivity to that probe.

### RV signal differentiation by Ab sample type

BM IgA (“maternal IgA”), cord blood IgG (“maternal IgG”), and 1-year infant IgG samples were assayed for reactivity against a total of 5,294 16-12mer peptides tiling RV A16 capsid proteins (VP1-4) and RV B52 and C11 complete polyproteins (**Fig. 2, supplemental data**). Every sample showed statistically significant reactivity to one or more probes for each RV genotype (**supplemental data**). Moreover, as evidenced by the heatmap banding patterns, there was notable similarity in antibody binding calls (**Fig. S2**) between different sample types and across all three RV species. When the data were realigned to directly compare analogous sequences, there was even stronger concordance in the most significant sites of epitope calls between A16, B52, and C11 (Pearson’s r 0.46 to 0.87), especially for the maternal IgG calls (**Fig. S3B**).

**Figure 2.**
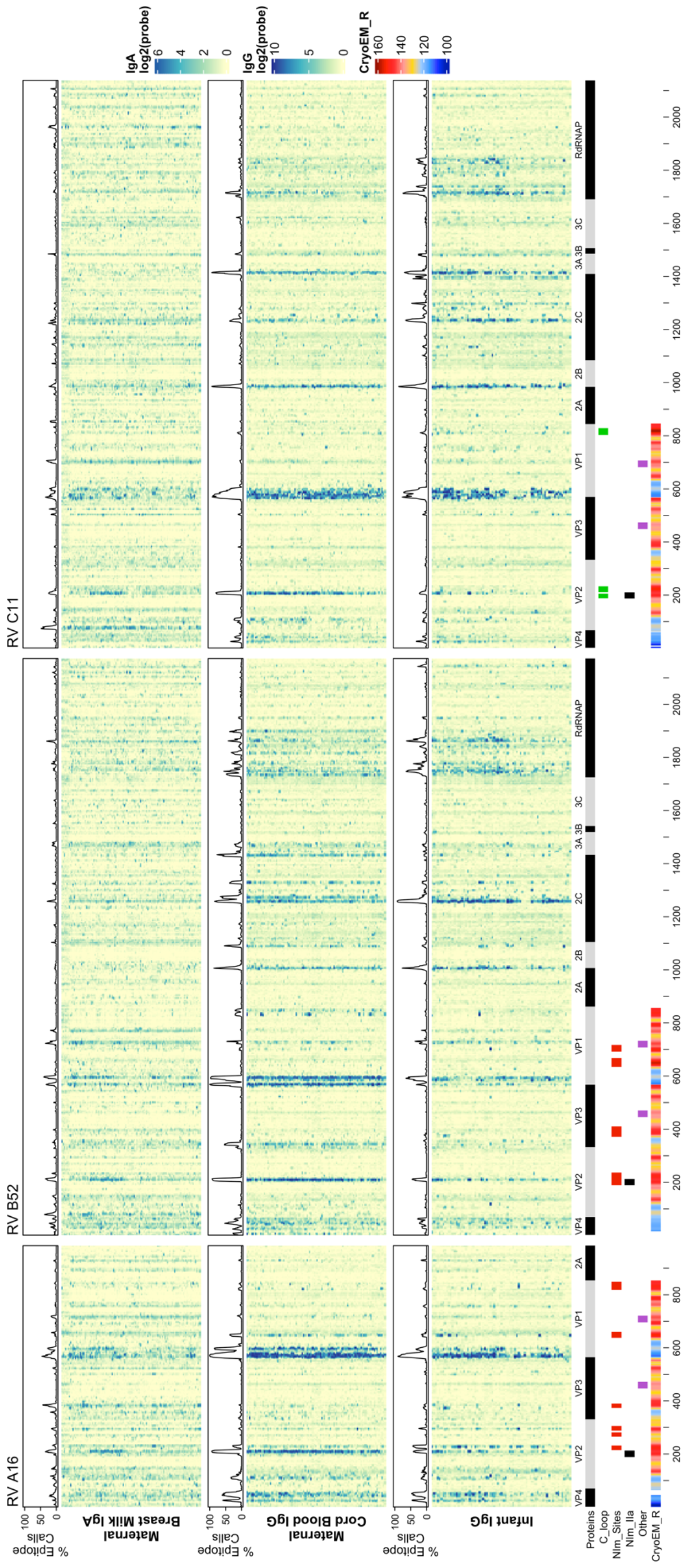
Maternal IgA and IgG and infant IgG reactivity to RV A, B, and C structural and non-structural proteins. Heatmap of antibody reactivity to tiled RV peptide sequences on a microarray. IgG and IgA sample data were processed as distinct populations. Above each sample population is a histogram of the percent of samples with statistically significant reactivity (i.e. epitopes) to that peptide. Several annotations are displayed below the heatmap including mature protein products, RV C C-loop residues (green), RV A2 and B14 NIm sites (red), B52 NIm-IIa sites-specifically (black), experimentally validated immunogenic peptides from Rossman et al. 1985 (purple), and cyro-EM derived radius values measuring the distance in Angstrom of a given capsid protein residue to the capsid structure center, for each probe the radius value is averaged over the residues of the peptide sequence, those probes with larger average radius values contain residues on the surface of the capsid structure.

It is also evident from the heatmaps that overall, and especially within the densest epitope regions, the maternal cord blood IgG epitope calls appeared qualitatively stronger compared with BM IgA. In addition, among mother-child dyads, there were more A16 and B52 epitope calls in cord blood compared to the infant samples at 1 year (**Fig. S3**). However, infant IgG samples had slightly more C11 calls compared to maternal IgG, possibly due to a greater frequency of RV-C infections among infants (Choi et al. 2021). Analysis of nasal mucus collected from infants during regular health checkups and self-reported illnesses indicate a significantly higher number of confirmed RV-C infections compared to RV-A (P = 0.0382; paired, one-sided Welch’s t-test) and RV-B (P = 1.092 x10-10; paired, one-sided Welch’s t-test), among tested infant samples.

Although IgG and IgA epitopes predominately co-localized, we identified 491 probes (from A16, B52, or C11) with disproportionate antibody binding between BM IgA and cord blood IgG samples (proportions test, FDR < 0.001). Most of these probes were bound by a greater proportion of BM IgA samples. Notably, a BM IgA-specific epitope was observed near the B52 and C11 VP2 N-terminus (spanning probes 76-83 and 72-83, respectively) and in C11 VP1 (probes 701-707) (**Fig. S4A-C**). Cord blood and infant IgG samples disproportionately bound 193 probes, with more infant samples binding to B52 and C11 non-structural protein sites than cord blood IgG. Several B52-specific sites were preferentially bound by cord blood IgG samples, including probes 332-349 (VP3 N-terminus), probes 826-839 (VP1) and 843-853 (VP1); there was little-to-no BM IgA or infant IgG recognition of these B52 epitopes.

### Reactivity with known surface neutralization sites

RV virions have 60 identical protein subunits arranged with icosahedral symmetry. Each subunit is derived from a single cleaved polyprotein precursor. The 3 largest proteins, VP1-3, contribute equivalently to the virion surface. A smaller fourth protein (VP4) is packed adjacent to the interior RNA (Rossmann et al. 1985). Neutralizing immunogenic sites (NIms), as defined by VP1-3 escape mutations to monoclonal antibodies, have been mapped in cell culture for A2 and B14 isolates. Exclusively confined to the outer virion surfaces, these reported locations generally tag flexible protein surface loops contributed by structurally adjacent sequences. The composite NIms for A2 (sites A, B, C) and for B14 (NIm-I, NIm-II, NIm-III) occupy roughly analogous surface topologies when the respective capsid coordinates are superimposed (**Fig 3**). The strong conservation among all RV structures, combined with homology-based sequence alignments allow prediction of comparable elements for equivalent viruses, especially within the same species. Each A2 NIm surface feature residue has direct homolog in the A16 proteome tiled on the peptide array; the B14 NIm locations, likewise, are predictors of analogous B52 sites (**Fig 2**, red bars in annotation).

**Figure 3.**
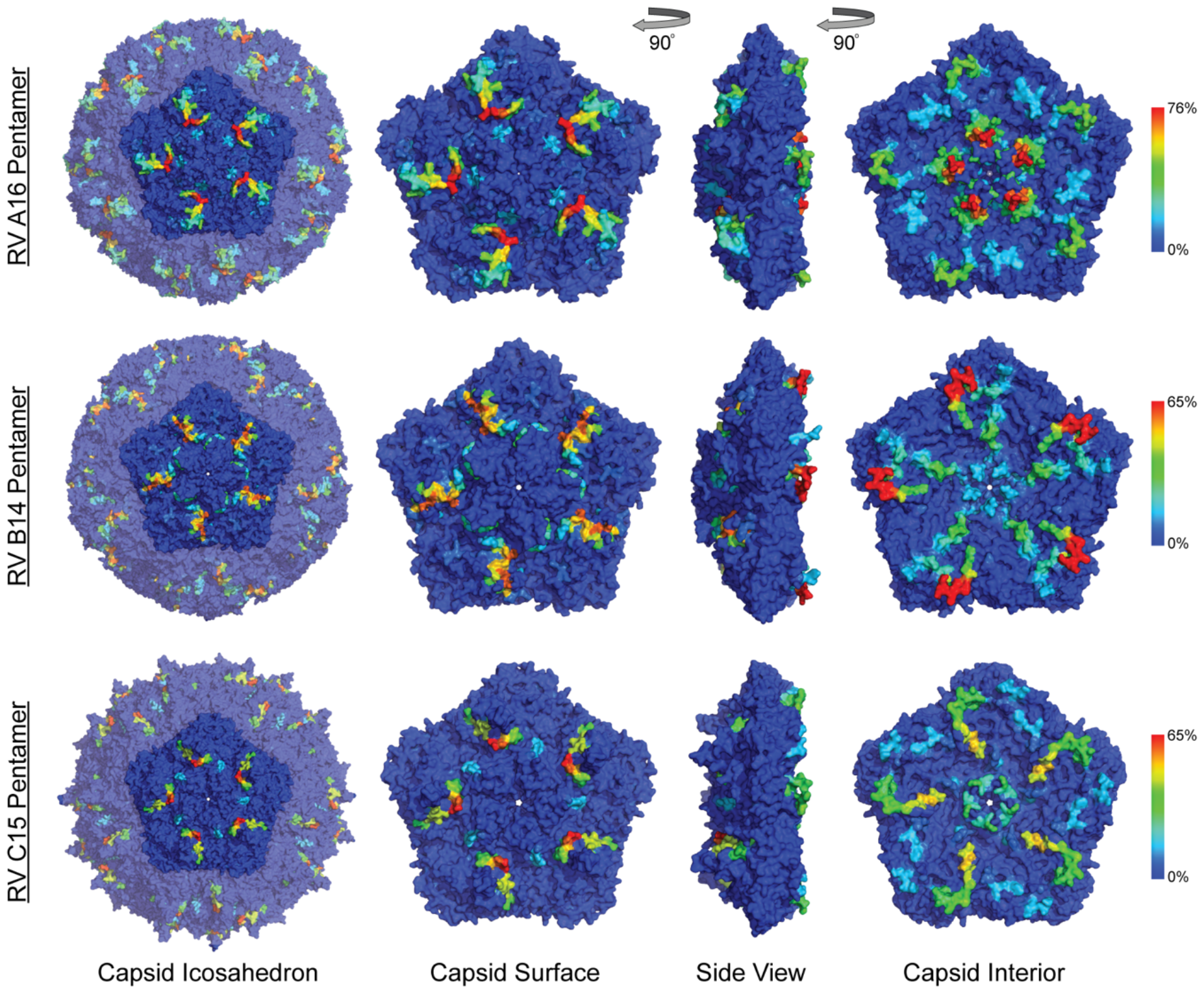
Anti-HRV antibodies to outer and inner surface of capsid proteins. Heatmaps of epitope calls as a percent of all samples for RV A16, B52, and C11 were overlaid on a coordinate file for A16 (1aym), B14 (4rhv) and C15 (5k0u) structures, respectively, available from the Protein Data Bank. Dark blue to red color scale represents the percent of all samples with statistically significant signal for the peptide sequence starting at each residue in the structure.

When the A2:A16, B14:B52 sequence and structure equivalents were annotated with observed array calls, reactivity to NIm sites was highly variable, dependent upon the source of the antibody and also on each protein’s residue contribution to that site (**Table 2**).

**Table 2.**
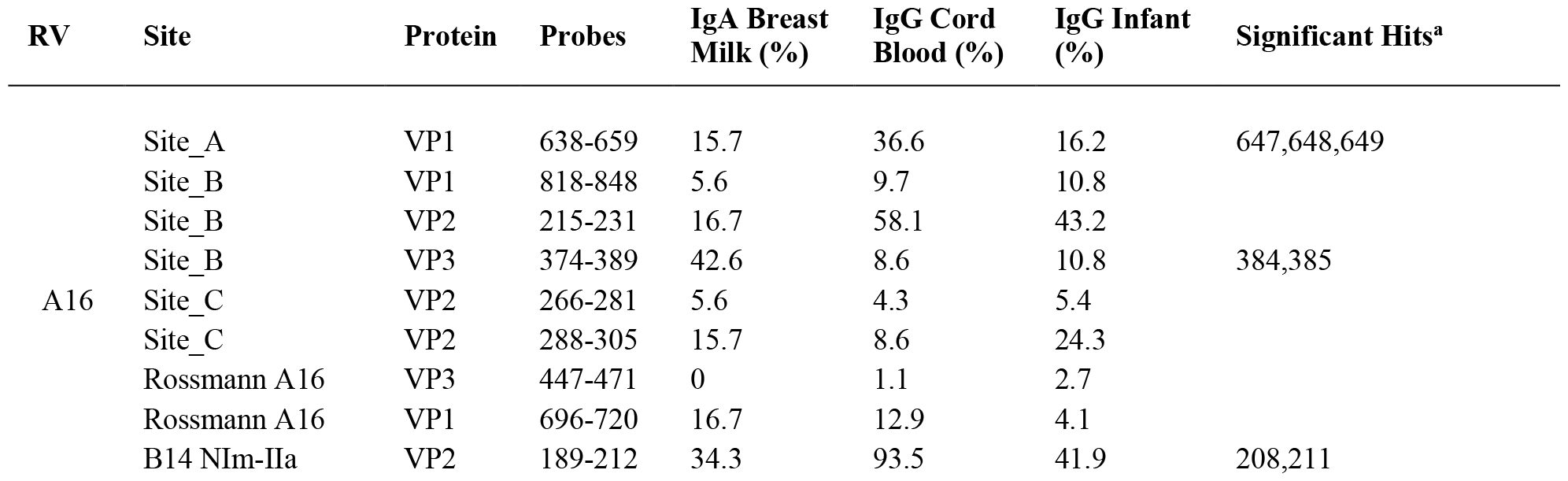
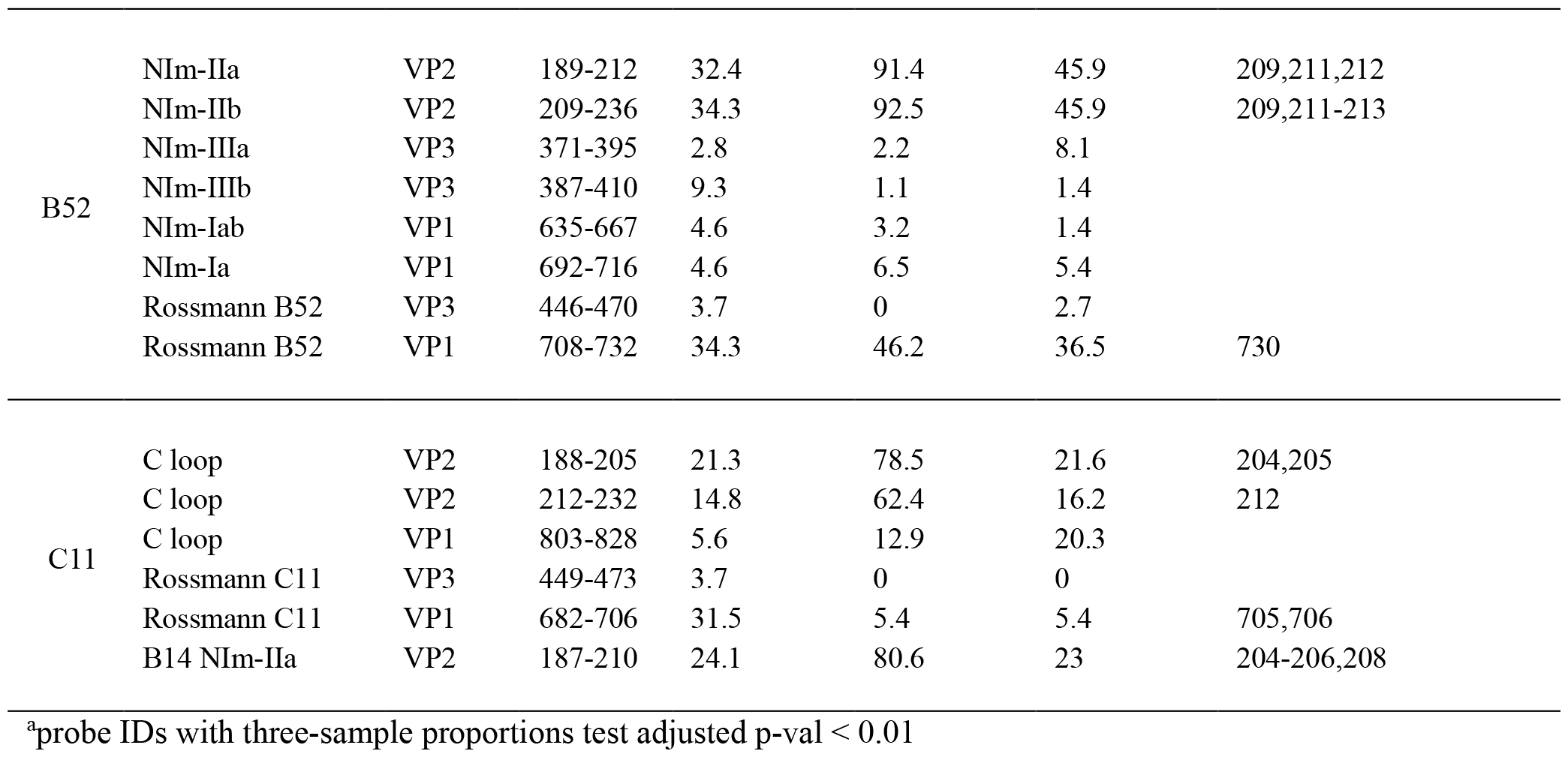

Reactivity to the annotated A2 NIm sites in A16, varied across sample types and antibody isotypes (**Table 2**). For example, reactivity to A2:A16 Site A was biased toward cord blood IgG. Here we define bias as a statistically significant difference in the proportion of sample binding. Reactivity to A2:A16 VP3 Site B was biased towards BM IgA. Reactivity to A2:A16 Site C, which includes two regions of VP2, was limited to the C-terminal residues of this NIm site, found in probes 288-305.

As many as 92% of cord blood IgG samples are bound to B14:B52 NIm-IIa and NIm-IIb. The B14 structure configures this region as a continuous VP2 surface sequence with slightly more external exposure than the analogous A16 Site B (VP2) segment. An orthologous VP2 epitope was also observed in A16 and C11 VP2 proteins (**Fig. 2**), which aligns to the B14 NIm-IIa site. Infant IgG reactivity values and epitope binding were highly correlated between the A16 and B52 B14 NIm-IIA site (**Fig. S5**). In A16 the B14 NIm-IIa site is distinct from and located just upstream of the A2 Site B NIm site and prominent A16 epitopes overlap each site. However, reactivity to NIm-IIa and Site B in A16 varied considerably depending on the sample type. While cord blood IgG and BM IgA primarily reacted to NIm-IIa, infant IgG reactivity varied considerably, with each sample reacting to either Site B or NIm-IIa but rarely to both. This finding raises the question of how much reactivity at this or any other site stems from putatively cross-reactive antibodies, accumulating with a higher likelihood in maternal samples.

In addition to NIm sites, several other capsid peptides have been validated experimentally in animal models as immunogenic, and possibly conserved as epitopes in related picornaviruses (Fig 2, purple bars in annotation). These include two B14 10mer capsid fragments in VP1 and VP2 (Rossmann et al. 1985; McCray and Werner 1987) (**Table 2**). The Rossmann VP1 peptide was bound consistently across all sample types and RV species. The B52 VP1 Rossmann peptide was bound by the greatest overall proportion of samples and was biased to IgG samples. In contrast, the C11 VP1 peptide was biased to BM IgA.

### Reactivity with the capsid interior

Aside from the known external surface virion neutralizing sites (NIms), multiple other capsid segments, in all three RV species, were bound by samples. Among the most prominent sites were peptides spanning the VP3 C-terminus and VP1 N-terminus. In mature virions, the majority of these residues are structurally interior, but when subunits are assembling, they are sequence-adjacent until separated by proteolytic cleavage. The cord blood and infant IgG samples show these regions to be highly bound (**Fig. 2**) consistently displaying as a twin pair of epitopes for A16 and B52, but somewhat more continuous in the analogous region of C11. BM IgA samples had the same patterns, but in fewer samples. Among infant IgG samples though, there was a great deal of preference variability for the upstream (VP3) or downstream (VP1) segment of the pair. The infant IgG samples differentially bound to 90% and 34% of the A16 upstream and downstream segments, respectively, possibly indicating that different physical antibodies bind to these two sites, or that they are not simultaneously exposed during antibody formation or considered contiguous for recognition.

### RV Non-Structural Protein Epitopes

RV nonstructural proteins are not encapsidated into virions, but they are present in the cytoplasm of infected cells and in airway secretions during illnesses. For completeness, the full proteomes of B52 and C11 were included on the peptide arrays. Several common reactive regions were found, most notably at the 2A protease C-terminus, the 2B N-terminus, the middle of 2C, and in the N-terminal half of the 3D polymerase. As with the capsid samples, reactivity was more pronounced for cord blood and infant IgG. Especially with the infant IgG samples, the C11 banding patterns (**Fig 2**) were more coherent and pronounced, and involved a greater portion of samples, perhaps indicating that certain discrete segments make especially viable epitopes within 2C and 3D, or even within 3A, 3B, and 3C.

### Reactivity with RV-C-specific VP1 surface loop

RV-C capsids are topologically distinct from the RV-A and RV-B and have never been mapped experimentally for neutralizing NIms. Species-specific deletions, clearly apparent in the aligned sequences, truncate several VP1 loops contributing to the five-fold region, and consequently remove the NIm-Ia and NIm-Ib homologs that would otherwise be immunogenic (Basta, Sgro, and Palmenberg 2014; Lau et al. 2009). Analogs to NIm-II and NIm-III are still surface exposed, but unique to each RV-C subunit, and centered within it, are a pair of species-specific adjacent exposed protein loops contributed by a VP1 C-terminal extension (residues 256-265 RV C15), and nearby VP2 residues (Liu et al. 2016). The VP1 portion of this “finger” or “C-loop” varies in length (16-23 residues), with up to 70% amino acid diversity (average 50%) among the 57 recognized RV-C types. The C-loop is hypothesized to function as a dominant neutralization site, an apparent structural substitute for the missing NIm-Iab (Liu et al. 2016). The contributing VP2 residues (136-138;160-165) at the base of the C-loop partially align to the B14 NIm-IIa site, further supporting the likely immunogenicity of this unique RV-C feature (**Fig. 2**).

Approximately 40% of all tested samples showed significant binding to C11 VP2 C-loop residues (**Table 2**), particularly in cord blood IgG samples. Only 6-20% of the combined samples showed reactivity to the more prominent C11 VP1 loop component. This was slightly more common among the infant IgG samples. Our accompanying viral diagnostics that followed these infants indicated only about 4% ever had a confirmed RV-C11 infection during their first year of life which is considerably fewer than those exhibiting reactivity to the C11 C-loop. While sparsity within our nasal swab PCR data is a likely contributor to this incongruity, it raised the possibility that RV cross-reactivity, at this or other prominent epitope sites, might contribute to the observed signals.

A potential test for cross-reactivity was embedded within our array design by the inclusion of 16mer peptides tiling VP1 C-loop from 57 RV-C and five putative RV-C genotype sequences (**Fig. 4**). The largest proportion of infant IgG samples (57%) bound to C33 and C17 VP1 C loop segments. In contrast, no tested infant samples bound to C42 (JF416320), even though as many as 24% of infant samples showed reactivity to a related C42 sequence (JQ994500). This pair of isolates differs by 6 (of 16) amino acids over the sampled region, a diversity that is typical for other RV-C pairs in the C-loop region.

**Figure 4.**
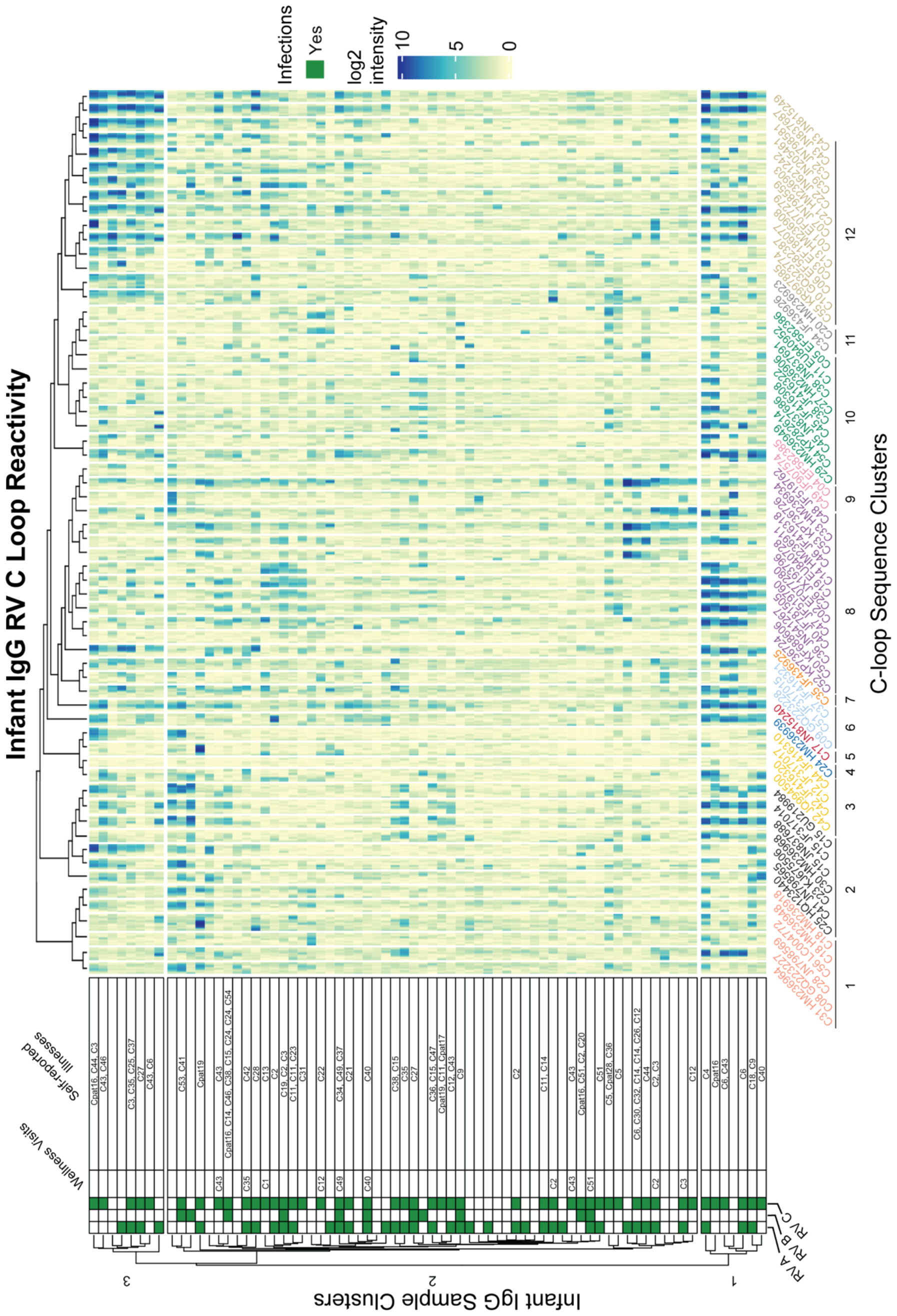
Infant IgG reactivity to HRV C-loops clustered by sample and C-loop sequence alignment. Heatmap of infant IgG (rows) reactivity to tiled HRV C-loop peptide sequences (columns) on a microarray. K-means clustering of infant IgG reactivity yielded three distinct groups with similar reactivity profiles among more homologus C-loop sequences as derived from multiple sequence alignments. A phylogenetic tree of 62 aligned HRV C-loop sequences was cut into 12 clusters yielding the grouping scheme shown here. C-loop sequence labels are concatenated from the RV type and Genbank accession number from which the sequence was obtained.

Since cross-reactivity likely depends on sequence diversity, the infant IgG data were reparsed according to a C-loop Neighbor-Joining tree. The phylogram closely resembled published RV-C trees based on complete VP1 or VP0 sequences (Simmonds et al. 2010; McIntyre, Knowles, and Simmonds 2013), and identified at least three groups of infant samples with distinct preference for homologous C-loop sequence clusters. Sample group 1 (N = 7) had predominant reactivity to C-loop cluster 12, sample group 3 (N = 8) had reactivity to subsets of C-loop clusters 8 and 12, while sample group 2 (N = 59) consisted of samples with limited or overall reduced C-loop reactivity. These results did not change when redundant probe sequences (i.e., probes common to more than 1 virus) were removed (data not shown) indicating that positive calls were not due to artefactually similar C-loop probe sequences. Within infant cluster 1, for example, there was an observed enrichment for C6 infections (P = 0.027; hypergeometric test), but no enrichment for other RV. Not only did several of these IgG samples bind the C6 probes they also bound those

C-loops most related to C6. As with the C11 data cited above, this again indicates the potential for cross-reactivity, at least for this epitope, within this phylogenetic group.

## Discussion

We profiled the IgG and IgA antibody binding landscapes of mother and infant samples against three rhinovirus species. This is the first proteome-wide examination of RV B-cell epitopes comparing maternal (cord, BM) transfer of IgG and IgA antibodies via placenta and breastmilk respectively, to the fetus and newborn, offering passive protection until infants develop active antibody responses to infection (1 year plasma). We also profiled reactivity to the prominent RV-C C-loop structures that were predicted to be highly immunogenic. Highly immunogenic epitopes displayed a striking degree of concordance across sample types, antibody isotypes, and RV types. RV- and antibody isotype -specific epitopes were identified for capsid and non-structural proteins. Many less common epitopes were also present (found in <10% of samples for a given sample type) that may represent private B cell receptor repertoires or possibly cross-reactivity from infection from heterotypic RVs.

By profiling cord and BM samples these experiments produced new information about the quantity and particular anti-RV antibodies that the placenta and breastmilk transfers to the infant in the form of IgG and secretory IgA. Breastfeeding reduces respiratory illnesses (Chonmaitree et al. 2016), but little is known about how BM antibodies protect against RV. One study observed that RV neutralization by BM was dependent on sIgA (May and Clarke 2000). For RSV, BM IgG neutralizing antibody titer was a strong correlate of immunity for infant RSV infections (Mazur et al. 2018). We observed an abundance of maternal RV epitopes common to both plasma IgG and BM IgA, including epitopes at known NIm sites, which may protect against RV infections in early infancy. We also observed BM IgA-specific epitopes on the virion interior and surface of B52 and C11.

RV NIm sites are structural antibody epitopes composed of residues brought in close proximity by protein folding and capsid assembly. Using our array composed of linear peptides, samples strongly reacted to array peptides spanning VP2, and to a lesser extent VP1 NIm sites. Others have also reported strong IgG reactivity to short linear peptides spanning the A16 NIm-II site (Sam Narean et al. 2019). The absence of reactivity at other NIm sites may indicate a greater dependence on the capsid tertiary and quaternary structure for antibody reactivity to these sites. NIm site reactivity may also require seroconversion from homotypic RV infection. The nearly ubiquitous reactivity to VP2 NIm sites suggests that these sites are not only less dependent on the capsid conformational structure but may also be highly cross-reactive.

Neutralization sites have not been characterized for RV-C. Our infant and maternal samples reacted to several C11 virion surface features, including the VP1 and VP2 flexible protein loop protrusion (“C-loop”). Examination of additional RV-C C-loop sequences demonstrated strong reactivity, especially from infant samples, and further investigation is warranted to determine if C-loops are RVC NIm sites. Despite the considerable sequence diversity of RV-C C-loops, we observed evidence of putative cross-reactivity at these sites bases on similar reactivity patterns between homologous C loops sequences. A recent study reported reciprocal cross-reactive neutralization between RV C2 and C40 in both mouse and human samples (Bochkov et al. 2023). Our findings also grouped together RV C2 and C47 C-loops as putative cross-reactive epitopes indicating that potential cross-reactivity patterns can be deconvoluted from antibody binding profiles collected from peptide microarrays.

Numerous IgG and IgA epitopes localized to the virion surface and interior and to non-structural proteins. These potentially novel surface epitopes may represent additional neutralization sites, non-neutralizing antibody-dependent protection sites that aid in viral clearance (Behzadi et al. 2020), or sites for intracellular inhibition of viral replication and assembly (Mazanec et al. 1992). One of the most striking and immunodominant epitopes observed, spanning the VP3 C-terminus and VP1 N-terminus, consists of residues largely internal to the RV capsid structure. A previous study also observed IgG1 and IgM reactivity to RV VP1 N-terminal peptides that do not contain surface exposed residues (Niespodziana et al. 2012). Interestingly, studies of poliovirus and other enteroviruses related to RV have also reported B cell epitopes within the VP1 N-terminus of that localize to the virion interior (G. K. Lewis and Chu-Pei 1992; Cello et al. 1993; Samuelson et al. 1994). This poliovirus N-terminal VP1 epitope has been characterized as a “T-B epitope pair” by which this B cell epitope also shows strong T cell reactivity thus leading to B cell activation. Several RV-A and C VP1 peptides can stimulate T-cell proliferation in humans suggesting that a VP1 immunogenic T-B epitope pair is conserved across enteroviruses (Samuelson et al. 1994; Gaido et al. 2016). For B-cell epitopes not residing on the virion surface, it is unclear what role they play in mediating the host immune response. Some have proposed this VP1 epitope may act to misdirect the host immune response towards non-neutralizing sites, thus aiding viral escape. How B-cell receptors encounter these “internal” epitopes is unclear. Possibilities include changes in capsid structure during host cell attachment (Fricks and Hogle 1990), spontaneous capsid conformational changes (aka “breathing”) (J. K. Lewis et al. 1998; Kolatkar et al. 1999), and viral protein release following cell lysis.

Our profiling of peptide-level antibody reactivity to three RV types strongly supports the existence of cross-reactive RV epitopes. Many past and recent studies have reported observations of RV cross-reactivity, which can be examined comprehensively using array technology. Given the abundance and diversity of RV types, high-throughput profiling of RV B-cell epitopes by peptide arrays, as demonstrated here, can identify pan-RV epitopes that may direct the development of improved RV diagnostics, vaccines, and therapeutics.

## Author Contributions

JMV: Writing – original draft with input from IMO, Data curation, Formal analysis, Software, Visualization, Investigation, Methodology; SJM: Writing – review & editing, Data curation, Formal analysis, Software, Methodology; SF: Performed breastmilk assays, Selected and prepared samples, Methodology, Writing – review & editing; AP: Created the array design, Data curation, Visualization, Investigation, Methodology, Writing – review & editing; YB: Writing – review & editing; HL: Performed peptide array assays, Writing – review & editing; RP: Implement array design and synthesis, Writing – review & editing; JT: Performed optimization assays, Methodology, Writing – review & editing; JG: Investigation, Writing – review & editing, funding acquisition; CS: Conceptualized the study, Provided and selected samples, Investigation, Writing – review & editing; IMO: Conceptualized the study, Created the array design, Writing – review & editing, Funding acquisition, Visualization, Investigation, Methodology, Project administration, Supervision.

## Non-standard abbreviations

BM: breast milk; NIm: neutralization immunogen site

## Acknowledgements

We thank the study families for their participation in this study. The authors are grateful to Dr. Deborah Chasman and Dr. Aubrey Barnard for their thoughtful comments and helpful discussions in preparing this manuscript.

## Data Availability Statement

All peptide array datasets and code used will be made available after publication.

