## Supplemental Figures for "Assessing Immune Factors in Maternal Milk and Paired Infant Plasma Antibody Binding to Human Rhinoviruses"

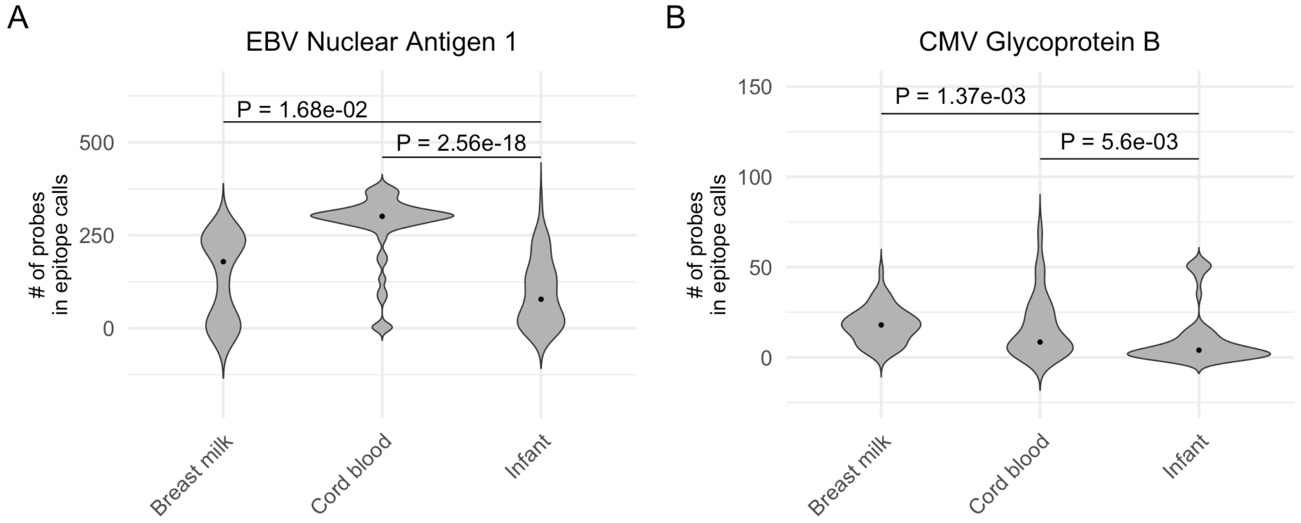


**Figure S1. Nascent Infant IgG Antibodies at 1 year Lower Compared to Maternal IgG.**
Violin plots showing the distribution of probes with statistically significant reactivity against control proteins (**A**) EBV nuclear antigen 1 and (**B**) CMV glycoprotein B among concordant mother-child dyads. Infants have fewer IgG epitope calls to these control proteins compared to their mothers (one-sided, paired t-test).

A

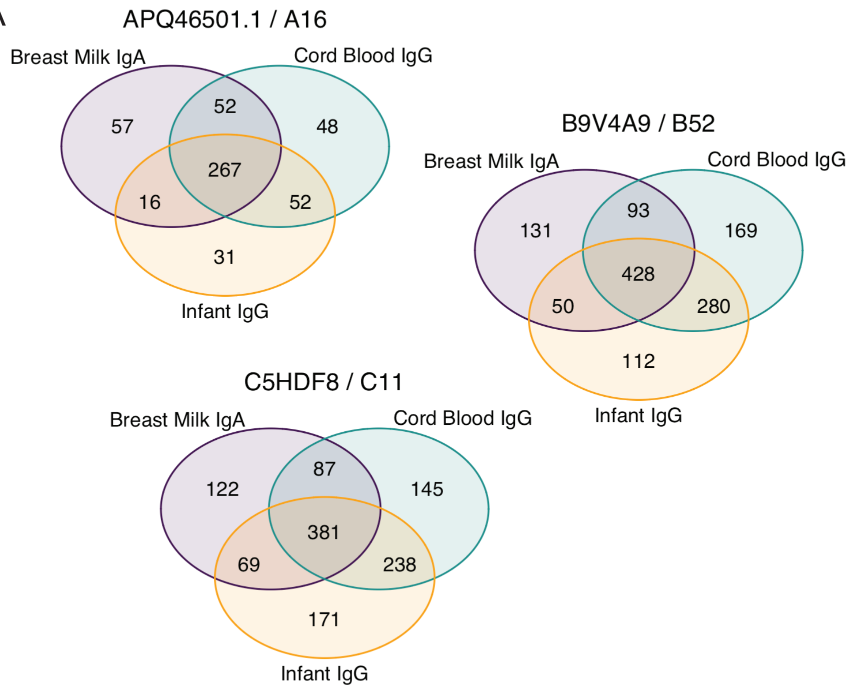


B

| | | A16 | B52 | C11 |
|---|---|---|---|---|
| **Breast Milk IgA** | **A16** | 1 | - | - |
| | **B52** | 0.61 | 1 | - |
| | **C11** | 0.57 | 0.72 | 1 |
| **Cord Blood IgG** | **A16** | 1 | - | - |
| | **B52** | 0.8 | 1 | - |
| | **C11** | 0.87 | 0.77 | 1 |
| **Infant IgG** | **A16** | 1 | - | - |
| | **B52** | 0.46 | 1 | - |
| | **C11** | 0.74 | 0.6 | 1 |

**Figure S2. High Degree of Epitope Concordance Between Sample Types and Serotypes.**
(**A**) Venn diagrams showing intersection of probes called as epitopes for each of the sample types for each HRV serotype. (**B**) Table of Pearson's correlation of aligned capsid protein sample epitope call histograms across the all HRV species in each sample type. Where correlation of k of N value of epitope calls for each probe was determined, with k as the number of samples with an epitope call for that probe and N = total number of samples.

A

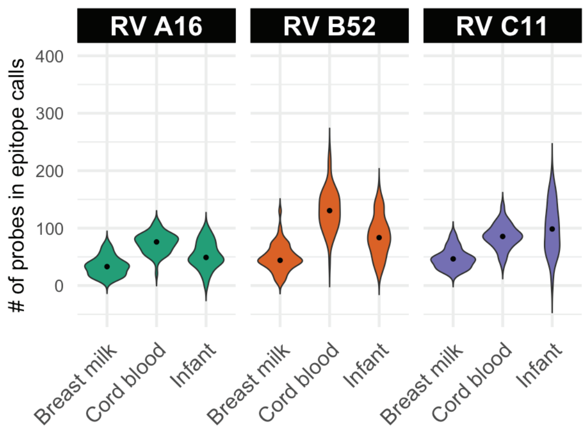


| | | Is < Cord blood IgG | Is < Infant IgG | Is > Infant IgG |
|---|---|---|---|---|
| **A16** | **Breast milk IgA** | 3.1e-19 | 5.1e-06 | n.s. |
| | **Cord blood IgG** | - | n.s. | 1.4e-06 |
| **B52** | **Breast milk IgA** | 7.4e-20 | 3.1e-08 | n.s. |
| | **Cord blood IgG** | - | n.s. | 3.8e-08 |
| **C11** | **Breast milk IgA** | 2.3e-14 | 2e-11 | n.s. |
| | **Cord blood IgG** | - | 2.9e-02 | n.s. |

B

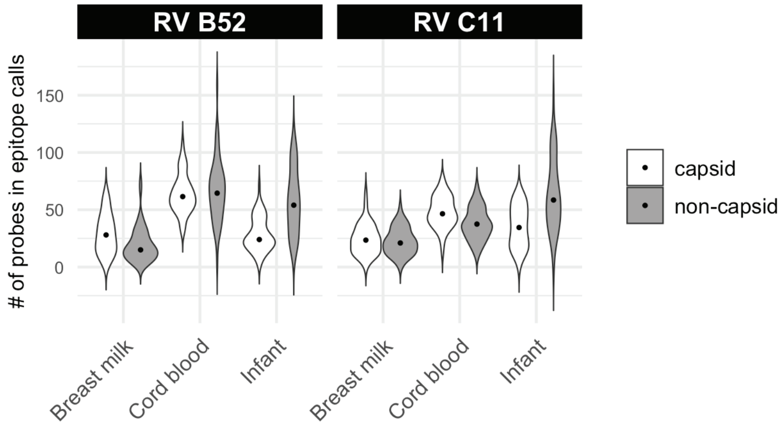


**Figure S3. Infant IgG Epitope Calls Decrease Compared to Maternal IgG and Increase Compared to Maternal IgA.**

(**A**) Violin plots showing the distribution of the number of probes with statistically significant reactivity (i.e. epitopes) to RV A16 (APQ46501.1), RV B52 (B94A9), and HRV C11 (C5HDF) among concordant mother-child dyad pairs. Except for C11, infant IgG samples show a statistically significant decrease in epitope calls compared to maternal IgG found in cord blood. Conversely, all infant IgG samples show a statistically significant increase in epitope calls compared to IgA from their mother's breast milk. Table reports p-values of paired, one-sided Welch's t test; n.s. = $p > 0.05$; "-" indicates tests was not performed. (**B**) Violin plots showing the distribution of the number of bound probes from RV B52 (B94A9), and HRV C11 (C5HDF), split by capsid and non-capsid proteins, among concordant mother-child dyad pairs.

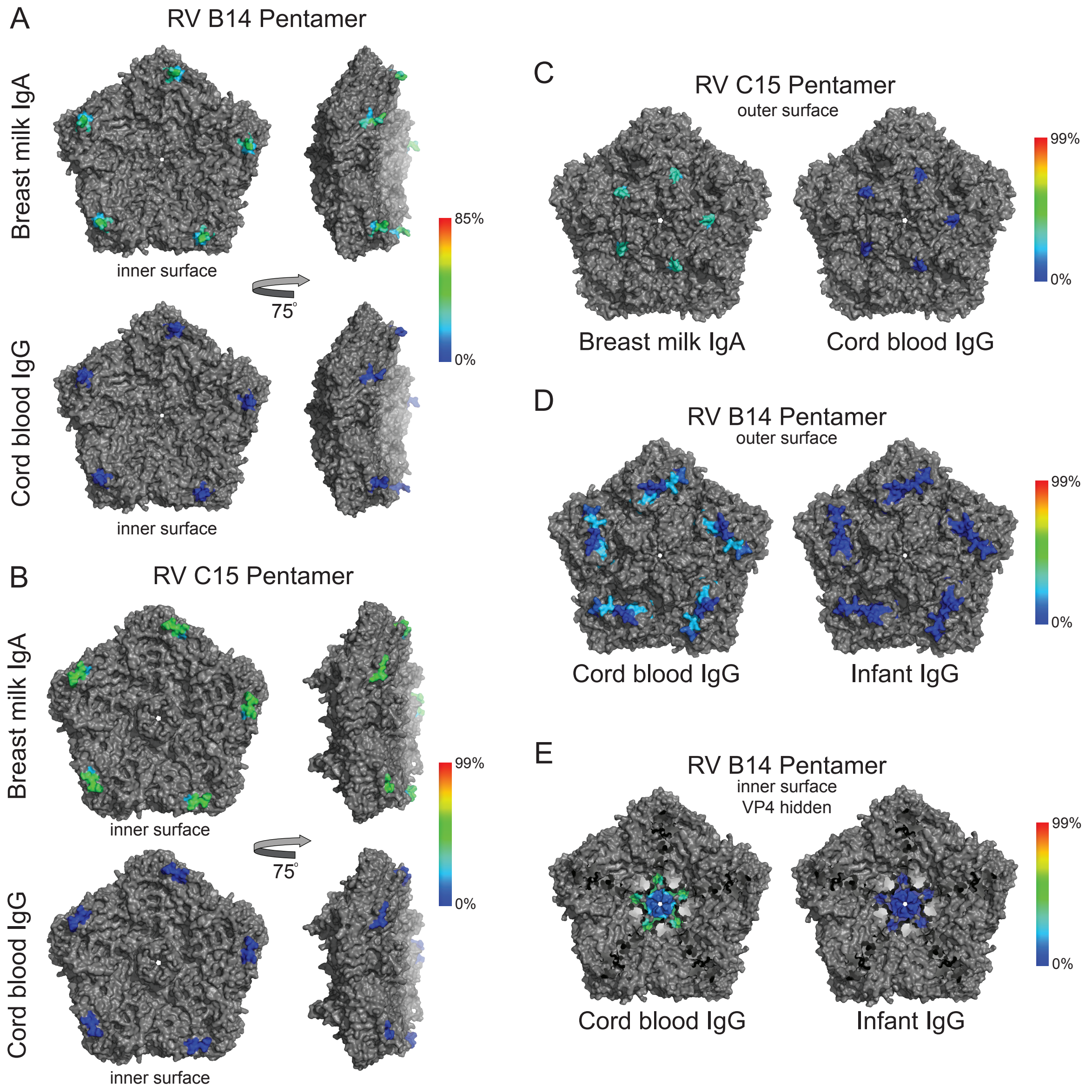


**Figure S4. B52 and C11 Antibody-Specific Capsid Epitopes.**
Heatmaps of epitope calls as a percent of BM and cord blood samples for RV B52 and C11 were overlaid on a coordinate file for B14 (4rhv) and C15 (5k0u) structures, respectively, available from the Protein Data Bank. A BM IgA-specific epitope at the VP2 N-terminus of both B52 (A) and C11 (B) along the virion interior. (C) A BM IgA- and C11-specific VP1 epitope along the outer surface. B52- and cord blood IgG-specific VP1 (D) and VP2 (E) epitopes. Dark blue to red color scale represents the percent test samples with statistically significant signal for the peptide sequence starting at each residue in the structure.

A

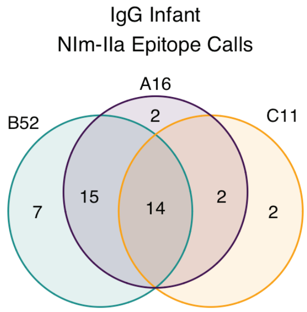


B

| | 1 | | | 2 | | | 3 | | | 4 | | | 5 | | | 6 | | | 7 | | |
|---|---|---|---|---|---|---|---|---|---|---|---|---|---|---|---|---|---|---|---|---|---|
| | A16 | B52 | C11 | A16 | B52 | C11 | A16 | B52 | C11 | A16 | B52 | C11 | A16 | B52 | C11 | A16 | B52 | C11 | A16 | B52 | C11 |
| A16 | 1 | 0.824 | 0.258 | 1 | 0.939 | 0.483 | 1 | 0.945 | 0.578 | 1 | 0.95 | 0.667 | 1 | 0.926 | 0.632 | 1 | 0.916 | 0.645 | 1 | 0.878 | 0.643 |
| B52 | 0.824 | 1 | 0.253 | 0.939 | 1 | 0.511 | 0.945 | 1 | 0.598 | 0.95 | 1 | 0.64 | 0.926 | 1 | 0.58 | 0.916 | 1 | 0.56 | 0.878 | 1 | 0.536 |
| C11 | 0.258 | 0.253 | 1 | 0.483 | 0.511 | 1 | 0.578 | 0.598 | 1 | 0.667 | 0.64 | 1 | 0.632 | 0.58 | 1 | 0.645 | 0.56 | 1 | 0.643 | 0.536 | 1 |

| | 8 | | | 9 | | | 10 | | | 11 | | | 12 | | | 13 | | |
|---|---|---|---|---|---|---|---|---|---|---|---|---|---|---|---|---|---|---|
| | A16 | B52 | C11 | A16 | B52 | C11 | A16 | B52 | C11 | A16 | B52 | C11 | A16 | B52 | C11 | A16 | B52 | C11 |
| A16 | 1 | 0.884 | 0.659 | 1 | 0.877 | 0.652 | 1 | 0.897 | 0.641 | 1 | 0.882 | 0.634 | 1 | 0.869 | 0.672 | 1 | 0.827 | 0.658 |
| B52 | 0.884 | 1 | 0.523 | 0.877 | 1 | 0.536 | 0.897 | 1 | 0.575 | 0.882 | 1 | 0.587 | 0.869 | 1 | 0.658 | 0.827 | 1 | 0.657 |
| C11 | 0.659 | 0.523 | 1 | 0.652 | 0.536 | 1 | 0.641 | 0.575 | 1 | 0.634 | 0.587 | 1 | 0.672 | 0.658 | 1 | 0.658 | 0.657 | 1 |

**Figure S5. NIm-IIa Cross Reactivity in HRV A, B, and C.**
(**A**) Venn diagram showing the intersection of infant IgG samples with NIm-IIa epitope calls in HRV A, B, and C. (**B**) Pearson’s correlation coefficient of infant IgG sample population probe intensity values for each of the 13 probes of the B52 infant IgG epitope spanning NIm-IIa.
